# Unravelling Form and Function: Improved function of engineered cardiac tissue through extra-cellular anisotropy

**DOI:** 10.1101/2021.08.06.455438

**Authors:** Jamie A. Cyr, Maria Colzani, Semih Bayraktar, Vera Graup, Richard Farndale, Sanjay Sinha, Serena M. Best, Ruth E. Cameron

## Abstract

Cardiac tissue engineering is a promising therapeutic option for myocardial repair after injury, however, so far engineered heart patches have shown limited translational utility due to poor electrical integration and tissue contractility. Emerging research suggests that scaffolds that recapitulate the three-dimensional structure of the native myocardium improve physiological function. Complex scaffold fabrication remains a technical challenge and the isolated impact of scaffold architecture on tissue function and cellular physiology is poorly understood. Here, we provided a direct comparison between isotropic and aligned collagen scaffold morphologies where all confounding physio-mechanical features, such as strut wall thickness and surface roughness are conserved. This enabled the independent and systematic assessment of the effects of pore macro-architecture on global tissue function and cellular maturation. We seeded our scaffolds with embryonic stem cell derived cardiomyocytes (hESC-CM) and measured tissue function through calcium signal transduction and dynamic contractile strain. The aligned tissue constructs facilitated improved signalling synchronicity and directional contractility. We further examined the influence of scaffold macrostructure on intercellular organization and intracellular development. Cells on aligned constructs conformed to the orientation of the scaffold macro-structure and were found to have phenotypic and genetic markers of increased maturity. Our results isolate the influence of scaffold macro-structure on engineered tissue function at multiple length scales. These findings inform the design of optimized cardiac tissue and expand the potential for engineered tissue in regenerative and model medical systems by reducing the gaps in tissue functionality that limit their utility.

## Introduction

Myocardial infarction (MI) results in permanent structural degradation of the ventricular wall; this leads to wall thinning and ventricular dilation that culminates in congestive heart failure^1–9^. Current treatments for MI, including the rapid limitation in the extent of infarction and blockage of maladaptive secondary pathways, can reduce cardiovascular morbidity and mortality. The remodelling process, however, still frequently leads to arrhythmias and decompensated heart failure contributing to cardiovascular morbidity and mortality without transplant ^10–12^. One possible alternative to conventional treatment protocols is the application of a cardiac patch which has been shown to encourage ventricular wall thickening, reduce cardiac wall stresses, and improve ventricular function^9,13–18^. Regenerative cardiac patches are engineered by populating biomaterial scaffolds with cardiomyocytes or other supportive cell types generally derived from stem cells ^11,19–27^. Scaffold materials commonly comprised either synthetic or natural polymers and are fabricated through a range of techniques including solvent casting^28^, gas foaming^29^, electrospinning^30^, rapid prototyping^31^ or freeze-drying^32^ where both the macro- and microarchitecture of the final scaffold is heavily dependent on the respective fabrication methodology ^33–37^. Clinical application of these techniques, however, remains limited by a lack of translational cellular functionality such as signalling synchronicity and appropriate physio-mechanical performance to promote cell survival, maturation and engraftment^37–40^.

Historically, translational designs for a regenerative cardiac patch have been structured by isotropic scaffolds or gels^41–43^. Recent improvements in scaffold fabrication technology have resulted in increasingly refined control over three dimensional (3D) scaffold architecture and subsequently the identification of biomimetic structure as an effective functional modulator ^17,30,44–47^. While these studies support structural anisotropy as a modulator of engineered cardiac tissue function, due to the complexity of scaffold fabrication and its interdependent relationship with both micro- and microarchitecture the isolated functional and developmental effects of scaffold structure are not fully understood.

In this work we aim to untangle the relationship between form and function in engineered cardiac tissue by isolating the functional and developmental effects of macro-architectural order. We provided a direct comparison between isotropic and aligned scaffold morphologies and systematically studied their effects on global tissue function and cellular maturation. To facilitate a direct comparison between scaffold morphologies we engineered a 3D ice-templated parent collagen (Devro Type 1, Bovine) scaffold with unidirectional alignment, from which we leveraged the inherent planar asymmetry to produce thin patches that were dominated by either isotropic or anisotropic pore architectures. Our use of a single scaffold fabrication technique ensures that all confounding physio-mechanical features, such as strut wall thickness, permeability and surface roughness were conserved across both conditions enabling the independent assessment of pore macro-architecture on tissue function.

Scaffolds seeded with human embryonic stem cell derived cardiomyocytes (hESCs-CM) were assessed for tissue function at a millimetre scale by characterizing time dependant contractile strain dynamics and both paced and spontaneous calcium signal transduction. We further executed detailed cellular analysis by measuring cell alignment, sarcomere length and gap junction formation. At a molecular level we evaluated differences in gene expression of cardiac maturation markers. Our results confirm the functional impact of scaffold architecture and further isolate the influence of scaffold macro-structure on engineered tissue function at multiple length scales. The field of cardiac tissue engineering has made rapid progress in recent years and this work will help expanding the potential for engineered tissue in regenerative and model medical systems by reducing the gaps in signalling capacity, contractility and phenotypic cellular maturity that have previously limited translational functionality and engraftment^9,30,38–40,48–51^.

## Results

### Generation of anisotropic and isotropic collagen scaffolds by directional ice-templating

Identification of the functional influence of scaffold structure is technically challenging due to the inherent link between manufacturing methodology and scaffold structure. To systematically study the isolated functional effects of architectural anisotropy in engineered heart tissue we decided to focus on three-dimensional collagen scaffolds generated by directional ice templating. Directionally freeze-cast scaffolds are characterized by an inherent structural asymmetry (Figure 1a) as shown by scanning electron microscopy (SEM) and micro computed tomography (μCT) that confirm different scaffold macrostructures across the transverse and longitudinal planes of the scaffolds. On the transverse plane the pore structure is composed of homogenous, isotropic, circular pores, while on the longitudinal plane the pore structure is composed of unidirectionally aligned pores (Figure 1 b-c). Fast Fourier transform analysis^52^ of each planar surface showed significantly increased alignment in the longitudinal plane (alignment order parameter: 0.658+/−0.084 and 0.175+/−0.056 normalized intensity units on transverse and longitudinal planes respectively) (Figure 1d). We exploited this feature to produce thin scaffold discs (8 mm diameter; 500-800 μm thickness) that were dominated by the architecture present on the circular face of the structure. Scaffolds sliced along the transverse plane of the parent scaffold were characterized by aligned pores (aligned scaffolds) whereas the ones sliced along longitudinal plane were mainly characterized by non-aligned circular pores (isotropic scaffolds). Due to the use of a common parent scaffold all other physio-mechanical properties such as pore size, strut wall thickness, permeability, interconnectivity, and surface roughness were maintained across scaffold conditions this allows us to systematically investigate the influence of structure on construct performance.

**Figure 1.**
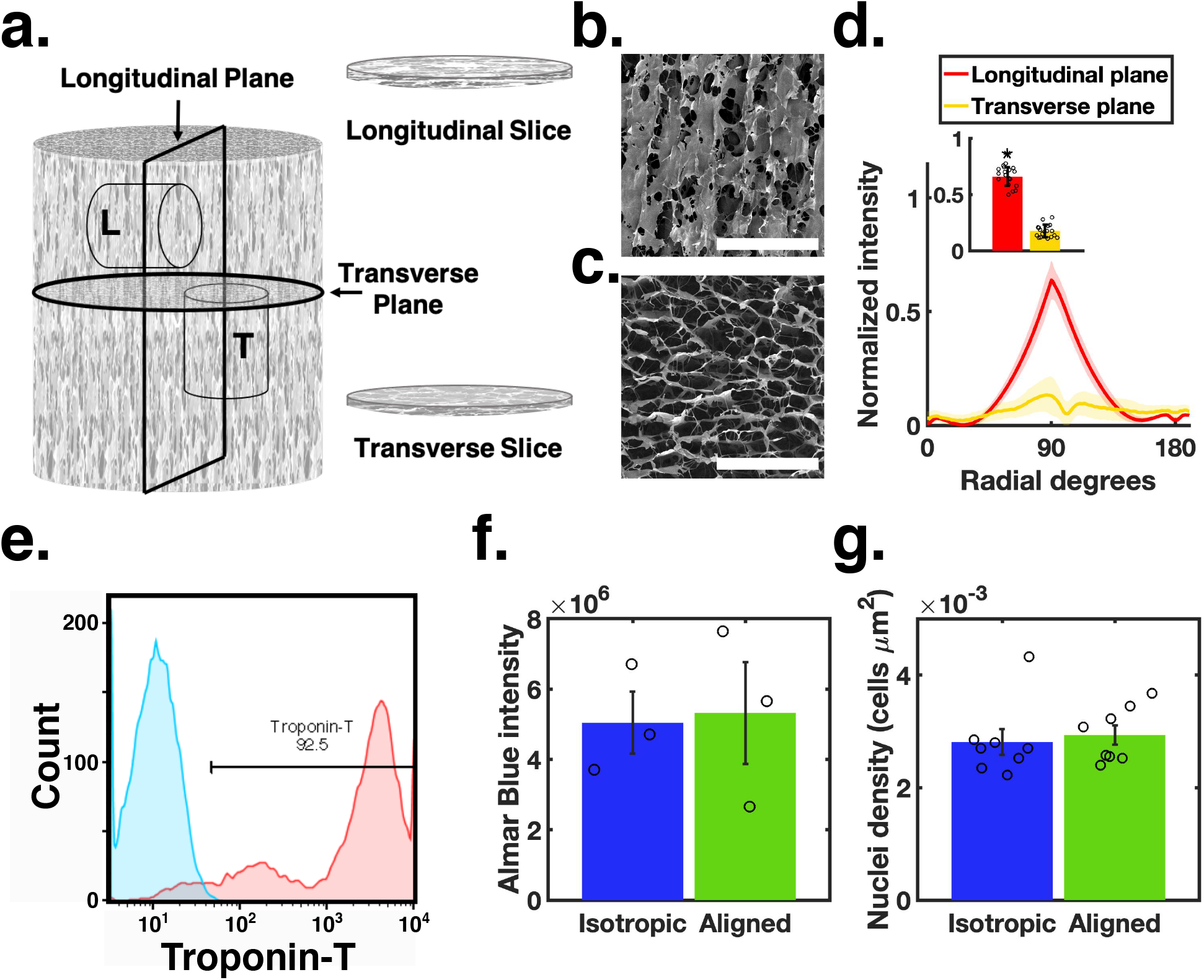
Engineered cardiac tissue **(a)** slicing schematic with biopsy punches and corresponding slices from the **L** longitudinal plane and **T** transverse plane **(b)-(c)** SEM micrographs of scaffold structure in the **(b)** longitudinal plane and the **(c)** transverse plane (scale bars 500 μm) **(d)** normalized FFT (f/f_0_) alignment at each radial orientation, insert shows the maximal alignment for each slicing plane (N=18) **(e)** a representative distribution of Troponin-T positive cells measured via flow cytometry. Blue indicates IgG antibody control; red indicates cells stained with an anti-Troponin-T antibody following lactate selection. **(f)-(g)** Engineered construct viability and cellular density measured through **(f)** Florescence intensity of Almar Blue (N=3) and **(g)** nuclei density for each construct architecture (N=8)

### Scaffold architecture does not affect cell viability and distribution

Once generated the scaffolds were populated with hESC-CM (over 90% Troponin-T positive Figure 1e) and cultured for 7 days. At day 7, cellular viability and density were assessed via Alamar Blue fluorescence viability assay and by calculating nucleus density on immunofluorescence images of cellularized scaffolds, respectively. Our results showed no significant difference in cell survival or cell distribution between the two scaffold architectures (Figure 1 f & g, 5.3×10^6^+/−2.5×10^6^ vs 5.0×10^6^+/−1.5×10^6^ arbitrary fluorescence units and 0.002+/−0.0004 vs 0.002+/− 0.0006 cells/μm^2^ on aligned and isotropic scaffolds respectively). We therefore assume that any observed functional differences are not due to preferential survival on a particular scaffold type.

### Scaffold anisotropy improves contractility

To analyse the contractility of the cellularized patches we utilised optical strain analysis to conduct a full field spatial-temporal assessment of construct deformation without constraining or disrupting the engineered tissue (Supplementary videos 1-2) ^46,53,54^. Scaffold architecture was found to dramatically influence deformation profiles and, thus, resultant principal strains (ε_1_ and ε_2_). Principal strain dynamics for both conditions occurred concurrently during contraction, however, isotropic constructs produced strains with equal and opposite magnitudes, indicating no net surface area change during the contraction (Figure 2a). The average maximal contractile strain (ε_1_) was - 0.018+/−0.008 with a maximal inotropic strain rate of ~0.1 s^−1^ and lusitropic strain rate of ~0.05 s^−1^ (Figure 2b). Spatial analysis of each principal strain at peak contraction further illustrates deficient contractile function as large variability in magnitude and direction was observed across the construct surface (Figure 2c & d). In contrast principal strains of aligned constructs indicated a net negative change in surface area during contraction, a finding more consistent with functional cardiac tissue^53^ (Figure 2e). The average maximal contractile strain magnitude for ε_1_ (0.145+/−0.039) was ten times greater than both the orthogonal component, ε_2_ (0.014+/−0.023), as well as the principal strains of isotropic constructs (0.018+/−0.008 and 0.015+/−0.001 for ε_1_ and ε_2_ respectively) (Figure 2 i-j). A similar relationship was observed in the strain rate profile (Figure 2f). The contractile function of aligned constructs was further confirmed through spatial analysis of peak contraction demonstrating coordinated full field principal strains during contraction (Figure 2g & h). Furthermore, the structural anisotropy was found to direct the deformation such that ε_1_ was oriented parallel to scaffold alignment (Figure 2h). Scaffold alignment dramatically impacted construct contractility such that aligned scaffolds facilitated increased contractility indicated by contractile principal strain magnitudes (Figure 2 i & j) and reduced directional variance (circular variance of principal angles of 0.158 +/− 0.097 for aligned and 0.395 +/− 0.098 for isotropic p=0.041) (Figure 2k).

**Figure 2:**
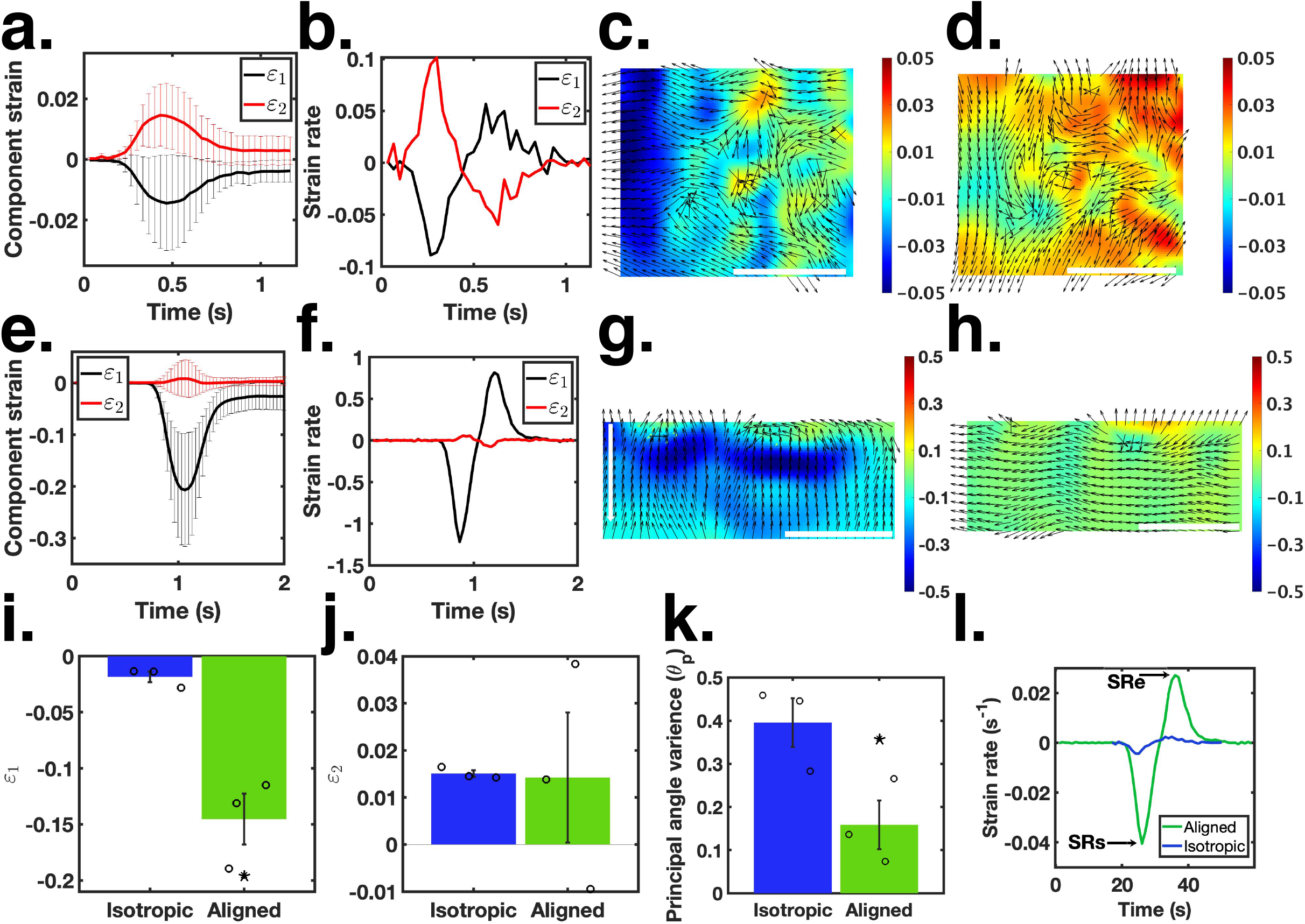
Live bright field imaging was performed on cardiomyocyte patches after 7 days of culture, video recordings were used to assess the construct deformation during contraction. Strain dynamics for isotropic **(a-d)** constructs and aligned constructs **(e-f)**. **a** & **e** Principal component strain in time ± standard deviation. **b** & **f** Mean strain rate for each principal component strain. **c** & **g** Spatial colour map for ε_1_ at maximum strain. **d** & **f** Spatial colour map for ε_2_ at maximum strain. **i** Maximum ε_1_ for all conditions **j** Maximum ε_2_ for all conditions. **k** variance of principal strain direction for all conditions. **l** Total construct strain rate for comparison with physiological strain rate. Isochrone scale bars are 1 mm; error bars represent standard error; N=3.

To understand how closely the anisotropic constructs resemble the native cardiac tissue, we next compared the dynamic strain profiles for each architectural condition with previously reported strain dynamics of *in vivo* myocardial deformation^53^. Three key characteristics of the strain rate profile have been identified: a global minimum during peak systole (SRs), a global maximum with reduced magnitude during diastole (SRe) and a small maximum during the isovolumetric contraction of diastole (SRa)^53^. The directional deformation dynamics produced by aligned constructs are consistent with the deformation profiles of native cardiac tissue *in vivo*, where ε_1_ is shown to be 2.5 times greater than ε_2_^53^. Similarly, with the exception of the intermediate peak due to isovolumetric contraction (SRa), the shape of the strain rate profile produced by aligned constructs is consistent with the strain rate produced by native myocardium. SRs and SRe were found to be −0.04 and 0.025, approximately 44% and 83% of physiologically recorded values respectively^53^ (Figure 2 l). Taken together, we demonstrated that aligned constructs not only enhance engineered construct contractility but also recapitulate the strain and strain rate characteristics of *in-vivo* cardiac tissue. These results are of particular relevance because the coordinated deformation between engineered and host tissue is paramount to ensure mechanical engraftment at the host/graft interface and has been one of the major functional limitations of engineered heart tissue^37,39,40^.

### Electrical connectivity is improved in anisotropic scaffolds

To explore the impact of long-range scaffold order on electrical conduction of cellularized constructs we utilize the analogous relationship between calcium transience and electrical impulse propagation by measuring spontaneous calcium activity (visualized by Fluo-4 AM) (Supplementary videos 3-4). Periodic fluorescence behaviour was observed for both structural conditions. Fast Fourier Transform analysis of each signal showed that engineered constructs with aligned architecture had a significantly faster pulse rate than the isotropic condition (0.55±0.09 Hz for aligned and 0.33±0.03 Hz isotropic p=0.019) (Figure 3a-c). We used the spatial distribution of spontaneous calcium pulse rates to quantify signalling continuity across the construct surface (Figure 3d-g). Isotropic constructs resulted in uneven pulse rate distributions with patchy heat-maps and high spatial variance (0.0174+/−0.012) across all samples. In contrast, aligned constructs resulted in largely uniform spatial pulse rate distributions and reproducibly small spatial variance (0.001+/−0.0006) (Figure 3h).

**Figure 3.**
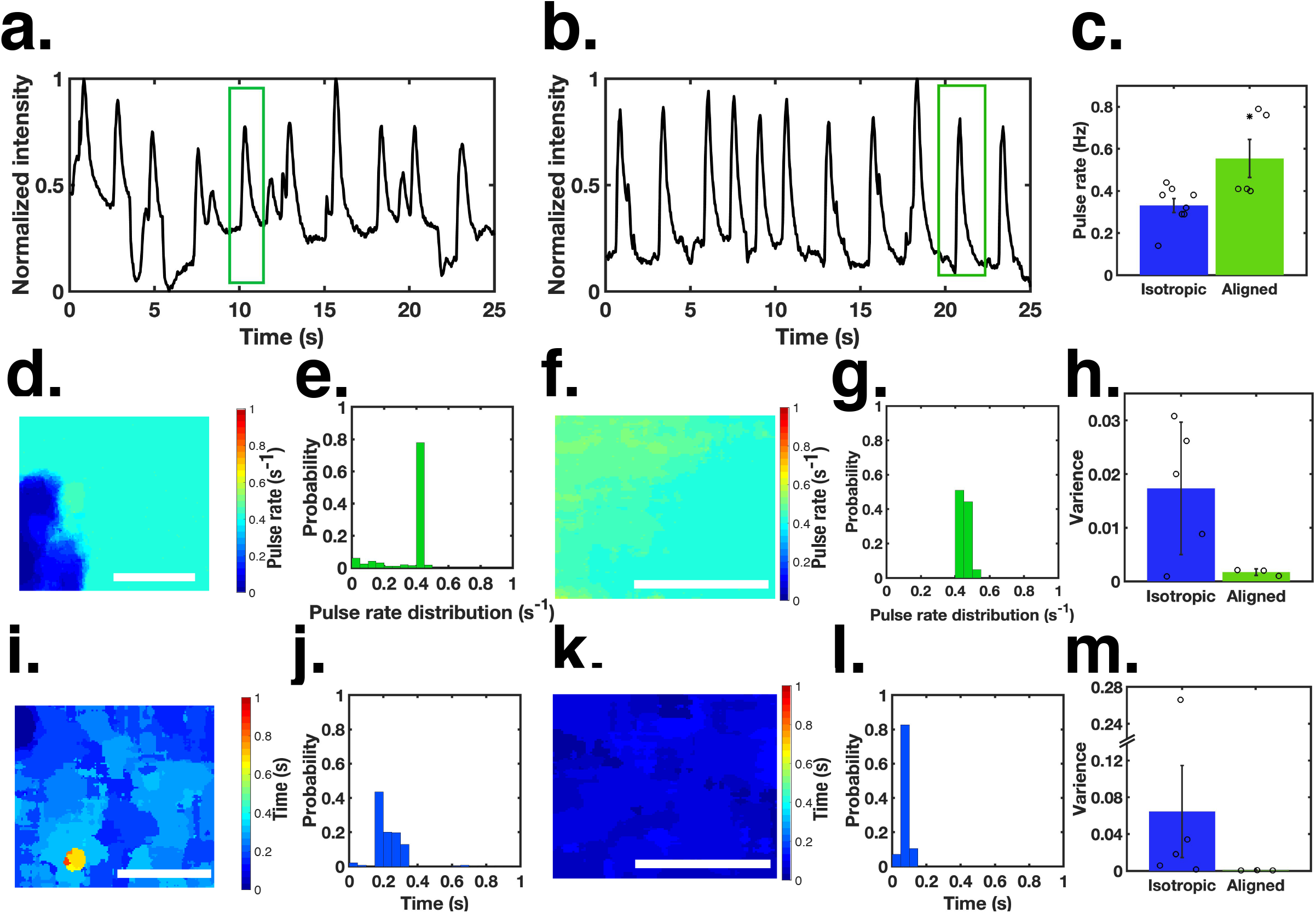
Live Fluo-4 AM calcium staining was performed on immature cardiomyocytes derived from H9 hESCs after 7 days of culture, video recordings of fluorescence dynamics were used to assess the temporal and spatial signalling uniformity **a** & **b** Mean fluorescence intensity in time for cardiomyocytes on isotropic and aligned scaffolds, respectively. **c** Pulse rate for all isotropic and aligned samples (aligned N=5; isotropic N=8). **d-g** Pulse rate in space and associated histogram for **(d)** & **(e)** isotropic constructs and **f** & **g** aligned constructs. **h** Spatial variance of the pulse rate for all isotropic (N=5) and aligned (N=3) constructs. **i-l** Time of peak fluorescence in space within a single pulse indicated by the green boxes in **a** & **b**, and associated histogram for **i** & **j** isotropic constructs and **k** & **l** aligned constructs. **m** Spatial variance of time of peak fluorescence within a pulse for all isotropic (N=5) and aligned (N=3) constructs; Scale bars represent 0.2 mm; error bars represent standard error.

The spatial-temporal coordination of peak calcium signalling was further analysed by mapping the spatial distribution of the time of peak florescence within a single pulse (Figure 3i-l). Here we show that isotropic constructs produced large intra-pulse spatial variance for the time of peak fluorescence (0.064+/−0.111), whereas aligned constructs resulted in predominantly concurrent signalling across the tissue surface (0.00053242+/− 0.00018947) (Figure 3m). The influence of scaffold architecture on spontaneous calcium signal conduction and coordination of engineered cardiac tissue further demonstrates the functional and translational utility of ordered extracellular macrostructure.

We next evaluated the calcium handling capacity of our constructs when subjected to electrical stimulation at 1 and 1.5 Hz. Calcium signalling on isotropic structures, irrespective of pacing frequency, displayed reduced regularity in calcium cycling when compared to anisotropic results (Figure 4 a-f). Additionally, while dynamic signalling on anisotropic constructs was able to conform to the external pacing frequency at both 1 and 1.5 Hz, isotropic constructs were not found to conform with a pacing frequency of 1.5 Hz as shown in figure 4f (1.473+/−0.092 vs 1.171+/− 0.215 Hz in aligned and isotropic scaffolds respectively p=0.089). We next characterized the calcium fluorescence wave form by the temporal characteristics illustrated in Figure 4g. Both time to peak and time to 90% decay of aligned constructs were shorter for both pacing frequencies when compared to those of isotropic constructs, as shown in Figure 4 h-k. The reduced temporal characteristics of the florescence waveform observed for aligned constructs indicate an increase in efficiency and responsivity in the calcium handling processes. Taken together, analysis of calcium dynamics demonstrates that anisotropic constructs facilitate improved signal transduction as well as intracellular calcium handling capacity, characteristics that have been shown to predict translational function of regenerative cardiac tissue^30,46,55–57^.

**Figure 4.**
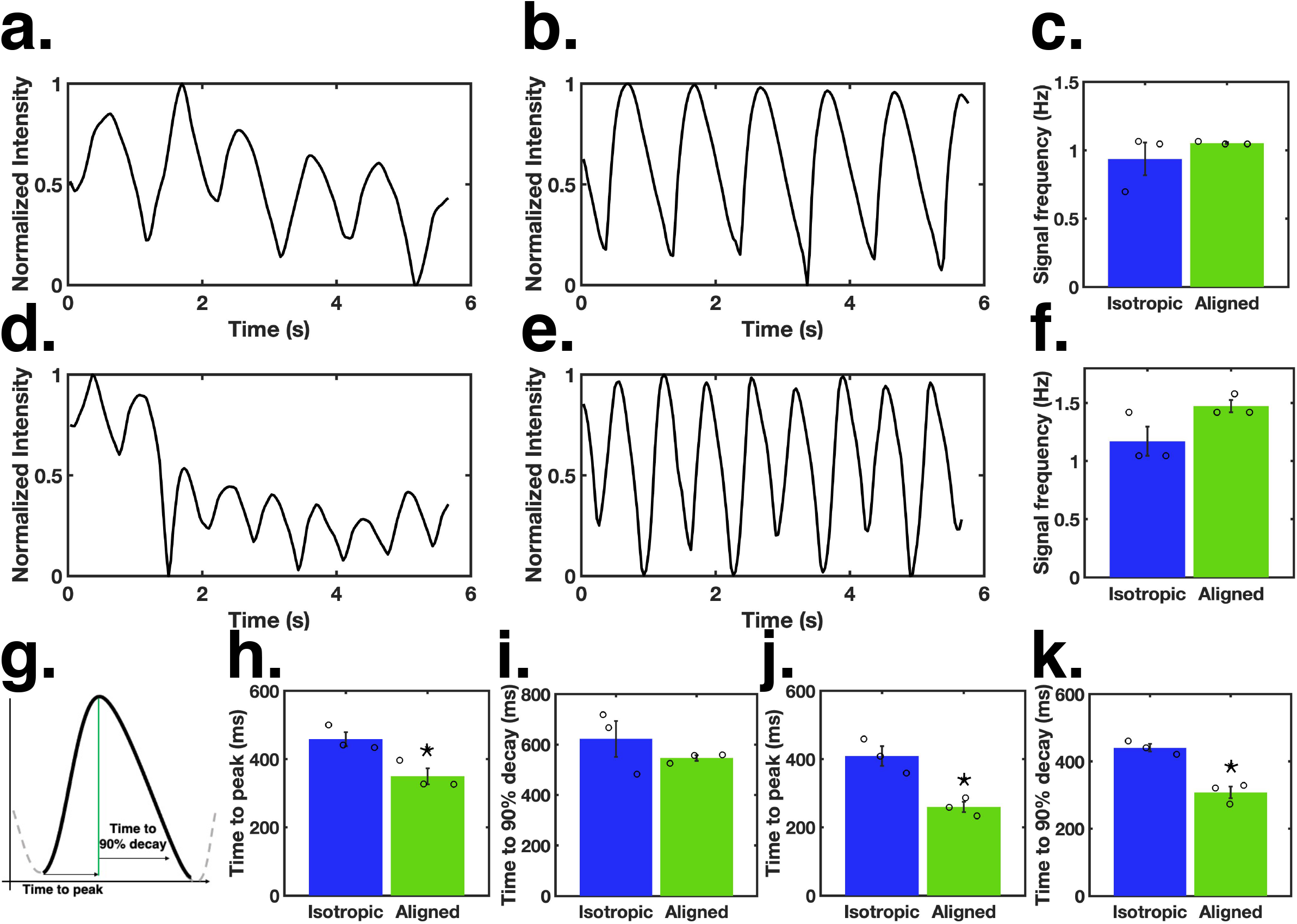
Paced calcium dynamics **a-b** normalized fluorescence intensity analysis with pacing at 1 Hz for (**a)** isotropic and **(b)** aligned constructs. **c** mean signal frequency for each structure. **d-e** normalized fluorescence intensity dynamics with pacing at 1.5 Hz for **(d)** isotropic and **(e)** aligned constructs. **f** mean signal frequency for each structure. **g** waveform diagram **h-k** waveform analysis of intensity dynamics for pacing at (**h**) & (**i**) 1 Hz and (**j)** & (**k**) 1.5 Hz; all error bars represent standard error; N=3.

### Scaffold anisotropy promotes cell alignment

To place these functional differences between scaffold architectures in context and further understand the impact of extracellular order on engineered tissue development we performed orientation and coherence measurements of seeded cardiomyocytes. Quantitative Fourier analysis of immunofluorescence micrographs of actin cytoskeletal structure (Phalloidin) was used to evaluate cellular orientation and intercellular alignment. Cells seeded onto isotropic scaffolds exhibited no preferential orientation direction (Figure 5 a-d), whereas the cellular population of aligned constructs had a more uniform orientation and conformed to the extracellular macrostructure (Figure 5 e-h) as demonstrated by increased cellular coherence (0.236+/−0.029 for aligned and 0.155+/−0.039 isotropic p=0.008) and reduced orientation variance (227.54+/−146.73 for aligned and 2.5×10^3^+/−1.6×10^3^ for isotropic p=0.012) (Figure i-k). Homogeneity of cellular directionality can help to explain the improved contractile performance of aligned constructs, as coordinated, directional cell shortening increases tissue level deformation^46,58–60^.

**Figure 5.**
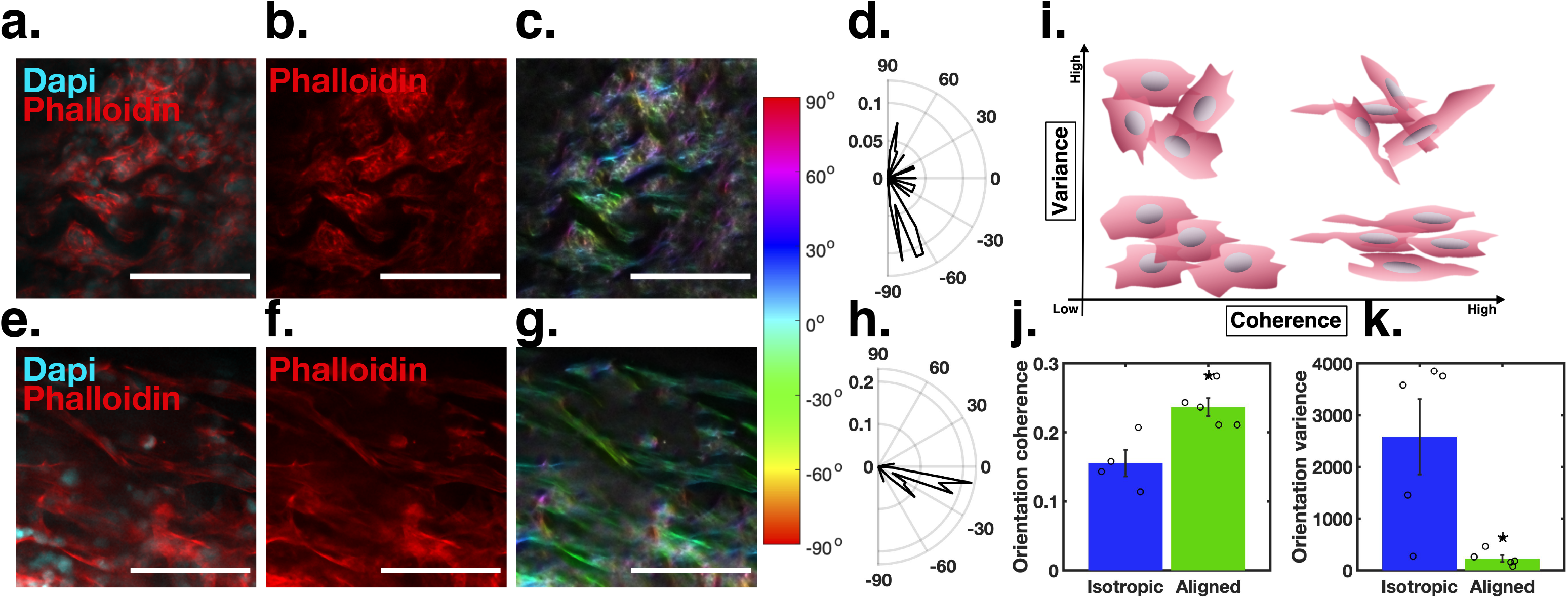
Cellular orientation of hESC-CM stained with phalloidin (red) and DAPI (blue) after 7 days on **(a)-(d)** isotropic scaffolds and **(e)-(h)** aligned scaffolds; scale bars represent 100 μm. **a** & **e** composite image. **b** & **f** Isolated Phalloidin channel showing actin organization. **c** & **g** actin orientation colormap resulting from Fourier transform orientation analysis over a moving pixel average of 2 pixels. **d** & **h** polar histograms of the actin orientation of cardiomyocytes within a single scaffold. **i** schematic of orientation variance and coherence measurements. **j** average actin orientation coherence for all isotropic and aligned samples. **k** average actin orientation variance for all isotropic and aligned samples; all error bars represent standard error; N=5.

### Scaffold anisotropy increases cardiomyocyte maturation

We next investigated in more detail the cardiomyocyte intracellular structural machinery, namely sarcomeres and gap junctions, to elucidate whether macrostructural tissue order influences phenotypic cellular development. We assessed sarcomere development by looking at sarcomeric α-Actinin staining. Cells seeded onto aligned scaffolds tended to have longer sarcomere subunits and displayed significantly increased banding prominence (Figure 6 a-f) indicating more advanced sarcomere development and cardiomyocyte maturation^61,62,66^. These results help to explain the improved contractility of aligned scaffolds shown in figure 2.

**Figure 6.**
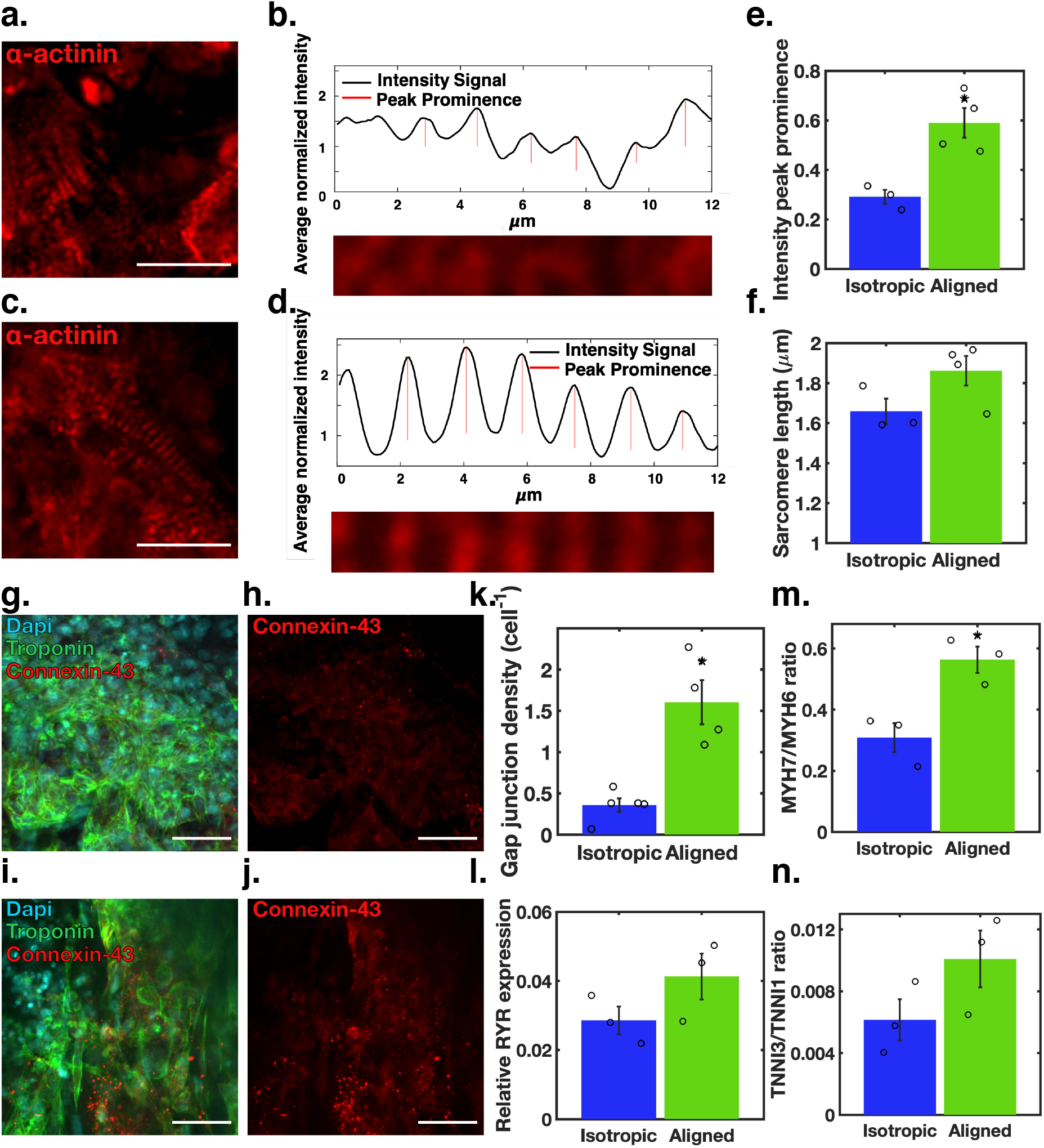
Intra-cellular structures and gene expression. **a-f** hESC-CM stained for sarcomeric α-actinin (red) after 7 days on **(a-b)** isotropic scaffolds and **(c-d)** aligned scaffolds; scale bars represent 20 μm. **b** & **d** representative quantification of sarcomere organization through relative intensity peak prominence along a single sarcomere chain. **e** relative intensity peak prominence (sarcomere intensity). **f** sarcomere length for cells on aligned (N=4) and isotropic (N=3) scaffolds. **g-k** hESC-CM stained for Dapi (blue) Troponin (Green) and Connexin (red) after 7 days on **(g-h)** isotropic scaffolds and **(i-j)** aligned scaffolds; scale bars represent 50 μm. **k** gap junction density for all isotropic (N=5) and aligned (N=4) samples. **l-m** qPCR quantification of relative expression of (**l**) RYR and **(m-n)** MYH7 to MYH6 and TTNI3 to TTNI1 expression ratios (N=3).

Staining for the gap junction protein Connexin-43 revealed that aligned structures facilitated a higher density of early gap junctions (1.603+/−0.534 gap junctions per cell) relative to isotropic structures (0.356+/−0.184 gap junctions per cell p=0.001) (Figure 6 g-k) supporting the increase electrical connectivity within the anisotropic patch previously observed with the calcium transient analysis.

Finally, we measured cardiomyocyte maturation by looking at expression of key maturation genes by qPCR. We observed greater ryanodine receptor (RYR2) expression in cells seeded onto aligned scaffolds when compared to those on isotropic structures (Figure 6l), possibly contributing to the shortened time to peak parameter of the calcium fluorescence wave form during pacing. The shift from expression of foetal type genes α-myosin (MYH6) and troponin-I 1 (TNNI1) to genes which have been linked to a more adult phenotype, β-myosin (MYH7) and troponin-I 3 (TNNI3) was also quantified. The MYH7/MYH6 and TNNI3/TNNI1 expression ratios are both increased in aligned constructs compared with the isotropic condition (Figure 6 m-n).

Taken together these observations support the notion that macro-structural alignment facilitates phenotypic maturation of hESC cardiomyocytes which in turn promotes the formation of engineered heart tissue that more closely recapitulates the native myocardium.

## Discussion

Clinical applications of regenerative cardiac tissue are limited in large part by an insufficient translational functionality, and lack of appropriate physio-mechanical cues to promote cardiogenesis, maturation and engraftment^9,38–40^. Currently, the electrical function and contractility of engineered cardiac tissues falls well below desired levels and is further hindered by inconsistent electrical integration with the host myocardium^37^. Here we demonstrated the functional impact of scaffold architectural order in cardiac tissue engineering by isolating and varying scaffold macrostructure while conserving all other biomechanical and material properties. We demonstrated that ordered scaffolds facilitated improved functional performance at multiple length scales: we assessed contractility and calcium signaling capacity at the tissue level, cellular orientation at the cellular level, gap junction and sarcomere development at the intracellular level and gene expression and maturation at the molecular level.

Contractility has emerged as an important determinant of functional cardiac tissues^45,46,63,64^. Conventional force-based methods of contractility measurement disrupt or limit the tissue structure. For example, Liu et al. 2020 utilized external grips to measure contractile force of neonatal mouse ventricular cardiomyocytes encapsulated in 3D methacrylated gelatin, the continuous unidirectional tensile resistance to tissue contraction imposed by the grips resulted in global tissue thinning between the constraints^45^. Recently, however, optical methods of measuring tissue contractility through strain analysis have been gaining traction. Shradhanjali et al. 2019 developed an Adaptive Reference-Digital image correlation (AR-Dic) method that enabled unbiased and accurate spatiotemporal deformation measurements of spontaneously contracting cardiomyocytes^63^. Similar methods have been utilised to characterize the spatiotemporal strain and strain rate dynamics of the native myocardium enabling direct comparison between engineered and native tissue dynamics^53,65^. We utilized an adaptive reference range methodology for digital image correlation to assess spatiotemporal deformation, strain, strain rate, and strain orientation during construct contraction. The presence of long-range structural alignment in engineered cardiac tissue facilitated not only increased deformation, but additionally a defined and directed synchronous contraction and effective tissue densification with coordinated surface area reduction. This behaviour closely matches directional deformation characteristics of *in-vivo* cardiac tissue that optimize cardiac output^53^.

Calcium handling dynamics have similarly been identified as an indicator of cardiac tissue function and are considered analogous to action potential propagation across the tissue^30,46,66,67^. We used spontaneous calcium florescence profiles to assess the electrochemical conduction capacity of cardiac constructs while paced calcium florescence waveforms indicated the cellular calcium cycling capacity of the populating cardiomyocytes. Aligned constructs facilitated spontaneous and periodic calcium signalling dynamics with a uniform spatial distribution as well as increased efficiency of calcium handling under paced conditions. In contrast isotropic constructs produced disjointed periodicity and uncoordinated spatial signalling dynamics with impaired responses to paced stimuli findings that have previously been associated with arrhythmia and myocardial damage ^30^. Taken together these results demonstrate that aligned structures facilitated an improved capacity to direct a synchronous electrochemical signal and conform to external pacing characteristics that are vital when assessing the functionality of regenerative cardiac tissue.

We further placed in context the observed functional differences between macro-structural morphologies by assessing cellular organization and phenotypic maturity. Unsurprisingly given the dramatic functional differences, it was found that the phenotypic development and intercellular organization were all positively influenced by increased architectural order of the engineered construct. The superior contractility of aligned constructs was supported by intercellular directional coordination of the cardiomyocyte population and by improved gap junction and sarcomere development^46^. Molecular analyses also suggested cellular and sarcomeric maturity as there was a shift from foetal sarcomeric proteins to the more mature forms. The improved calcium signalling dynamics and implied electrical conductivity were also supported by findings at multiple length scales. At the intracellular level the increased gap junction development observed in aligned constructs will improve intercellular signal propagation and electrical conductance. At a molecular level the increased expression of the ryanodine receptor helps to explain the increased responsivity and intracellular calcium handling efficiency and further validate a more mature phenotype. While these results clearly support our claim that macro-structural extracellular order alone can improve engineered cardiac tissue function, it is difficult to ponder a direct mechanism by which long-range alignment influences gene expression when local surface and mechanical characteristics at a micro- and nano- scale are conserved. We instead propose an indirect mechanism by which phenotypic cellular maturity is enhanced.

Bouchard and colleagues found that the application of increasing external electromechanical stimulation, gradually, hastened the maturation process of pluripotent stem cell derived-CM^64^. We hypothesise that long range anisotropic architecture facilitates a more coordinated contractile behaviour via globally organized cellular orientation. This enhanced and uniform contraction is effectively an auto-loading system, in which the amplified contractile force, facilitated by structural alignment, serves to cyclically stimulate the tissue construct and enhance the rate of cellular maturation^64^. Therefore as maturity increases, so too does the magnitude of the external stimulation which, in turn, further encourages maturity, creating an autologous loop that parallels the work done by Bouchard et al. 2019^64^. The effect is significantly reduced for the isotropic structure, as no long-range cellular order is present.

Several approaches have been developed to produce 3D engineered cardiac tissue, however, recently the focus has shift towards the study of macroscopic scaffold architecture and its effects on cellular phenotype^17,45,68^. Our study demonstrates, through a direct and systematic comparison, that anisotropic extracellular structure enhances the functional biomimetic capacity of engineered heart tissue at multiple length scales. Despite the necessity for further study to fully deconvolute the relationship between tissue structure and cellular maturation, our work shows how the application of macro-scale structural order directs engineered tissues towards a phenotype resembling the native myocardium and informs the design of an optimal tissue engineered myocardium for regenerative medicine and disease modelling applications.

## Methods

### Scaffold production

#### Collagen slurry preparation

A 1 w.t.% suspension of insoluble type I bovine dermal collagen (Devro) was prepared in 0.05 M acetic acid solution (Sigma-Aldrich UK). The mixture was left at 4°C to swell for 24 hours and homogenized in a blender at 22,000 rpms for 6 minutes. Gas was removed from the solution using a vacuum chamber (VirTis SP Scientific Wizard 2.0); the pressure was ramped from 750 torr to 2000 mtorr in 10 minutes. The slurry was allowed to habituate to room temperature (25°C).

#### Directional Ice-templating

Collagen slurry (9 mL) was loaded into a cylindrical polycarbonate mould (30 mm height, 20 mm internal diameter, 40 mm external diameter) with a copper base (2 mm thickness). The mould was placed onto a PID temperature controlled cold finger cooled with liquid nitrogen and programmed to hold at −10°C for 1 minute followed by cooling at a rate of 0.2°C min^−1^. The top of the mould was exposed to the ambient environment.

After solidification, moulds were dried in a freeze drier (VirTis SP Scientific Wizard 2.0) at 0°C under a vacuum of less than 100 mtorr for 20 hours.

#### Cross linking

Cross-linking was carried out using a ratio of 5:2:20 EDC:NHS:COOH groups in collagen was used to cross link at 5% of the standard (5:2:1)^69,70^. Cross-linking reagents were dissolved in 95% ethanol and scaffolds were soaked for 2 hours within the mould. Scaffolds were washed (5 × 5 min) with deionised water.

After cross-linking scaffolds were freeze dried (VirTis SP Scientific Wizard 2.0) with a cooling rate of 0.2°C min^−1^ to a primary freezing temperature of −20°C. Drying occurred at 0°C under a vacuum of less than 100 mtorr for 20 hours.

#### Slicing

An 8 mm biopsy punch and sliced with a straight razor to a thickness of 500-700 mm. Aligned structures were cut such that the circular face of the scaffold was parallel to the longitudinal plane of structural alignment. isotropic scaffolds were cut such that the circular face of the scaffold was parallel to the transverse plane of structural alignment as shown in Figure \ref{fig:MethodsScaffoldSlice}.

### Scaffold Imaging

Scanning electron microscopy (SEM) micrographs were taken of scaffolds prior to cross linking. Collagen scaffolds were-sputter coated with gold for 2 min at a current of 20 mA. All micrographs were taken using a JEOL 820 SEM, with a tungsten source, operated at 10 kV.

X-ray micro-computed tomography (mCT) images (Skyscan 1172) were taken of each scaffold with a voltage of 25 kV, current of 138 mA and a pixel size of 5.46 mm. Reconstructions of mCT images were performed with NRecon software by Skyscan.

### Scaffold Analysis

Reconstructions were divided into nine volumes of interest (2.5 × 2.5 × 6.5 mm^3^) dispersed across the bottom, middle, and top of the structure. Pore size analysis was applied to each transverse slice within the regions of interest. ImageJ software was used to binarize and watershed transverse slices and particle analysis was employed to compile pore size data. The pore sizes were analysed and visualized in MATLAB R2018a.

Fast Fourier Transform analysis was used to assess pore alignment according to the method laid out by Ayres et. al 2008^52^. 2D fast Fourier transform analysis was performed and radial sums of the resultant transform were collected in ImageJ. Pixel intensity for each radial direction was normalised by the minimum and plotted in MATLAB R2020a. The degree of preferential alignment was termed e_AOP_ and utilised to compare between samples.

#### Cell Differentiation

CH9 hESCs were maintained and differentiated according to protocols described by Iyer et al. 2015^71^ and Mendjan et al 2014^72^. An overview of the procedure is as follows: H9 hESCs were seeded into 12 well plates coated with Matrigel (Corning) filled with CDM-BSA supplemented with ROCK inhibitor (Millipore, 1 mm) at a density of 10^5^ cells cm^−2^. Media was changed after 3 hours to CMD-BSA, supplemented with FGF-2 (20 ng ml^−1^), Activin-A (50 ng ml^−1^), BMP-4 (10 ng ml^−1^, R&D) and LY294002 (10 mM Tocris). Media was changed again after 42 hours to CDM-BSA supplemented with FGF-2 (8 ng ml^−1^), BMP-4 (20 ng ml^−1^), Retinoic Acid (SIMGA, 1 mM), endo-IWRI (1 mM, TOCRIS). Cells were maintained with this media, refreshed every 48 hours, for 4 days. Media was changed to CDM-BSA supplemented with FGF-2 (8 ng ml^−1^), BMP-4 (20 ng ml^−1^) for an additional 2 days. Media was then changed to CDM-BSA with no cytokines and replaced every 48 hours. Spontaneous beating is generally observed 8-10 days after seeding.

#### Cardiac Cell Selection

Differentiated cardiac cells were metabolically selected via lactate selection. Media was removed from beating cardiomyocytes on 14 day. The wells were washed with PBS and TryPLE (Life technologies) (500 ml per well in a 12 well plate) was added. Plates were incubated at 37°C for 8-12 minutes, until dissociated. CDM BSA and DNase (DNase I Solution (1 mg ml^−1^) cat. 7900 Stemcell Technologies) diluted to 1:500 stock (1 mg ml^−1^) was added, 1 ml per well. Cells were collected in a falcon tube and centrifuged (3 minutes at 1200 rpm). Cells were resuspended in CDM BSA to a concentration of 1×10^6^ cell ml^−1^. Rock inhibitor (Y-27632 cat. 11573560 Millipore) (1:1000) was added. Cells were plated in a 6 well plate (2×10^6^ cells well^−1^) and incubate at 37°C for 8-12 hours. Media was removed from wells and lactate media (DMEM without Glucose/pyruvate with non-essential amino acid (cat.554084 Gibco) (1:100 from stock solution) and Sodium lactate (cat. L7022-10G SIGMA) (1:250 from 1M stock,4 mM final concentration)) was added (2 ml well^−1^). Cells were incubated in lactate media for 96 hours, media was refreshed after 48 hours.

The lactate selected hESC-derived cardiomyocytes were pelleted via centrifugation and resuspended in Fixation/Solubilization solution (BD Cytofix/Cytoperm Fixation/Permeabilization Kit, Biosciences) for 20 mins at 4°C. Cells were then pelleted by centrifugation and resuspended in 1x BD Perm/Wash Buffer containing anti-Cardiac Troponin-T APC antibody or Isotype control (Miltenyi Biotech) and incubated for 2 hours at 4°C. Cells were washed three times in 1x BD Perm/Wash Buffer, and then resuspended in phosphate buffered saline containing 0.1% BSA and 2 mM EDTA. Data was acquired on BD LSRFortessa™ Flow Cytometer and analysed with FlowJo™ v9.

### Scaffold Cellularization

Scaffolds described in section 0 were sterilised in 70% EtOH for 30 min. The EtOH was removed by PBS washing 3×5 minutes prior to scaffold conditioning with cell culture media (CDM BSA) for 1 hour in preparation for cell seeding. Cardiomyocytes were dissociated using TrypLE (Life technologies) and seeded at a density of 2 × 10^6^ cells per scaffold in CDM BSA supplemented with ROCK inhibitor 1 μM.

#### Analysis of Construct Performance

##### Viability

PrestoBlue Cell Viability Reagent (Thermo Scientific) was added to culture media according to the manufacturer’s instructions after 7 days of culture. Cells were incubated with the dye for 4 hours. Media was then sampled and fluorescence at 560 nm was analysed using VICTOR Multilabel Plate Reader (Perkin Elmer). Media containing PrestoBlue incubated in empty wells was used as background control.

##### Strain Analysis

Bright field videos were recorded on an Axiovert inverted microscope (Zeiss) using a Sony LEGRIA camera. Strain analysis was performed on bright field video samples with Ncorr digital image correlation software run on MatlabR2020a. The scaffold structure under bright field provided a reliable speckle pattern with sufficient contrast for analysis. A subset radius of 30 pixels and spacing of 5 pixels was used with the high strain option enabled. To avoid error due to global translation, the reference image was redefined after each beat, while the scaffold was in a relaxed state. Principal strain calculations were performed with MatlabR2020a. Principal angle characterization was performed using circular analysis of diametrically bimodal circular distributions^73^.

##### Calcium Dynamic Analysis

At day 7 after seeding, Fluo-4 AM (10 μg ml^−1^, Life technologies) was added to the cell culture media for 30 min at 37°C.Scaffolds were then transferred in Tyrode’s buffer and videos were recorded either with no stimulation or while pacing at frequency of 1 and 1.5 Hz using c-PACE EM pace (IONOPTIX). Videos were recorded on an Axiovert inverted microscope (Zeiss) using a Sony LEGRIA camera.

Video analysis was performed in MatlabR2020a. Fluorescence intensity was normalized and mean intensity was plotted against time. Both intensity peak frequencies and Fast Fourier Transform analysis were used to calculate pulse rate. For samples that did not exhibit spatial deformation during calcium fluorescence, pulse rates were also calculated for each pixel to indicate global signalling uniformity. Individual pulse times were recorded for each pixel and the temporal signalling uniformity in space was visualised through isochrones in MatlabR2020a.

##### Immunocytochemistry

Cell-seeded constructs were washed once in PBS then fixed for 1 hour with 4% PFA. The cells were subsequently permeabilised with 0.1% Triton (Sigma), 0.5% BSA (Sigma) in PBS for 15 min before blocking with 3% BSA (Sigma) in PBS for 1 hour. Incubation with primary antibody (diluted accordingly) was then performed. Constructs were then washed in PBS and incubated overnight with the appropriate secondary antibody, or phalloidin where appropriate, overnight. Constructs were then washed and stained with DAPI (Sigma, 1 μg ml^−1^) for 1 hour prior to imaging. Micrographs were obtained using an SP-5 confocal microscope (LEICA). Primary (I) and secondary (II) antibodies are listed in Table 1.

**Table 1.** Primary and secondary antibodies

| *Antibody* | *Supplier* | *Cat. Number* |
| --- | --- | --- |
| *mouse monoclonal anti-Cardiac Troponin T (I)* | Abcam | Ab8295 |
| *Rabbit polyclonal Anti-Connexin 43 (I)* | Abcam | Ab11370 |
| *Rabbit polyclonal Anti-Sarcomeric α-actinin (I)* | Abcam | Ab68167 |
| *Phalloidin - 555* | Abcam | Ab68167 |
| *Chicken anti-mouse 488 (II)* | Life Technologies | A-21200 |
| *Donkey anti-rabbit 568 (II)* | Life Technologies | A-10042 |

##### Cell density

Dapi stained nuclei were counted with particle analysis in ImageJ. The cell density for 200 μm^2^ regions of interest was calculated in MatlabR2020a for each scaffold.

##### Cellular Alignment

F-actin staining was used to characterise cellular spreading and cytoskeletal alignment. The F-actin orientation and coherence of cardiomyocytes after 7 days of culture was measured for 50 μm^2^ sections (27 measurements were taken per scaffold) with the OrientationJ plugin for ImageJ. The intra-scaffold variance was calculated for each individual scaffold.

##### Sarcomere development

Individual sarcomere chains were isolated from confocal images showing a-actinin such that the sarcomere band spanned the height of the region of interest. Banding intensity was characterize^61,62^. Fluorescence intensity was normalised by the minimum fluorescence (*f_0_*) such that, *f_norm_=(f/f_0_)-1*. The mean fluorescence intensity signal was plotted along the length of the sarcomere chain and the relative prominence of each intensity peak was measured in MatLabR2020a to calculate sarcomere intensity. Sarcomere width was defined as the signal wavelength.

##### Gap junction density

Immunofluorescence staining of Connexin-43 was used to visualise gap junction structures through fluorescence microscopy. Mature gap junctions were counted with particle analysis in ImageJ. The gap junction density per nucleus was calculated in MatlabR2020a for each scaffold.

##### RNA extraction, retrotranscription and RT-qPCR

RNA was extracted using GenElut Mammalian Total RNA Miniprep Kit (Sigma) according to the manufacturer’s instructions. RNA (100 ng) was subsequently retrotranscribed to complementary DNA (cDNA) using Maxima First Strand cDNA Synthesis Kit (Thermo scientific). RT-qPCR was performed using Fast SYBR Green Master Mix on a 7500 Real-Time PCR System using GAPDH as a housekeeping gene. All primers were designed to span an intron-exon junction, and are listed in Table 2. The relative expression of mRNA was obtained using the DCt method.

**Table 2.**
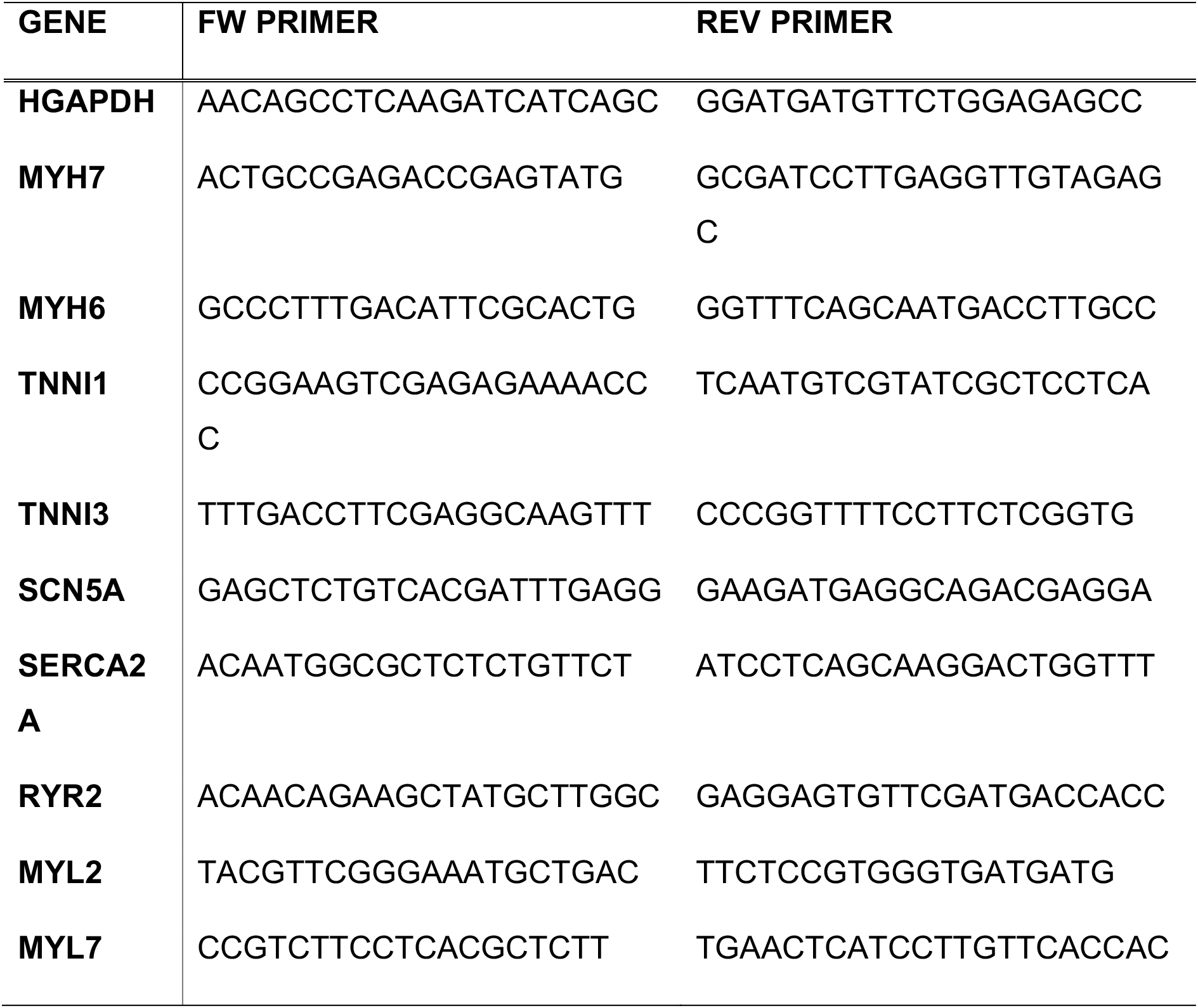
RNA Primers

##### Statistics

Experiments were executed three times in triplicate. A t-test with 95% confidence interval was used to determine statistical significance. Error bars represent standard error throughout. N-numbers are reported throughout

## Acknowledgements

The authors would like to thank the Wellcome Trust-MRC stem cell institute imaging facility and the NIHR BRC Cell Phenotyping Hub.

## Author contributions

JC and MC designed and performed the experiment, analysed the data and edited the manuscript. SB helped with the calcium transient analysis. VG performed preliminary experiments that informed this work. RF held a grant that supported this work. RC, SB and SS designed, conceived the study and interpreted the data and edited the manuscript. All authors approved the manuscript.

## Funding

This work was supported by the British Heart Foundation Oxbridge Centre for Regenerative Medicine RM/17/2/33380 and a BHF Senior Fellowship FS/18/46/33663 (S.S.). S.S. was also supported by the British Heart Foundation Centre for Cardiovascular Research Excellence. M.C. was supported by CRM (RM/17/2/33380) and also has support from BHF grant SP/15/7/31561. J.C. was supported by the Gates Cambridge Fellowship. S.B. and R.C. were supported by EPSRC Established Career Fellowship EP/N019938/1. V.G. received support from a PhD studentship from the BHF Cambridge Centre of Excellence. We also acknowledge core support from the Wellcome Trust and MRC to the Wellcome Trust – Medical Research Council Cambridge Stem Cell Institute. This research was funded in whole, or in part, by the Wellcome Trust [Grant Number: 203151/Z/16/Z]. For the purpose of Open Access, the author has applied a CC BY public copyright license to any Author Accepted Manuscript version arising from this submission.

## Open data

The original data from this paper is available at doi XXX.XXX.XXXX (site to be added)

## Notes

### Competing Interest Statement

The authors have declared no competing interest.

